# Single-cell analysis reveals different age-related somatic mutation profiles between stem and differentiated cells in human liver

**DOI:** 10.1101/547893

**Authors:** Kristina Brazhnik, Shixiang Sun, Omar Alani, Milan Kinkhabwala, Allan W. Wolkoff, Alexander Y. Maslov, Xiao Dong, Jan Vijg

**Author notes:** These authors contributed equally to this work (Kristina Brazhnik, Shixiang Sun). Correspondence to: K.B. and J.V.

## Abstract

Accumulating somatic mutations have been implicated in age-related cellular degeneration and death. Because of their random nature and low abundance, somatic mutations are difficult to detect except in single cells or clonal lineages. Here we show that in single hepatocytes from human liver, an organ normally exposed to high levels of genotoxic stress, somatic mutation frequencies are high and increase substantially with age. Significantly lower mutation frequencies were observed in liver stem cells and organoids derived from them. These results could explain the increased age-related incidence of liver disease in humans and stress the importance of stem cells in maintaining genome integrity.

---

Genome integrity is critically important for cellular function. Evidence has accumulated that loss of genome integrity and the increasingly frequent appearance of various forms of genome instability, from chromosomal aneuploidy to base substitution mutations, is a hallmark of aging (*1, 2*). However, somatic mutations occur de novo across cells and tissues and are of low abundance. Hence, thus far, of all mutation types only chromosomal alterations could readily be studied directly during in vivo aging using cytogenetic methods (*3*). More recently it has become possible to also study low-abundant somatic mutations by whole genome sequencing (WGS) of single cells or clones derived from single cells (*4–7*). Such studies confirmed that like chromosomal alterations also base substitution mutations accumulate with age in human brain (*8, 9*), intestine and liver (*4*).

Due to its high metabolic and detoxification activity liver is more likely to be adversely affected by various damaging agents than most other organs. In humans this may cause age-related loss of function, most notably a severe reduction in metabolic capacity and multiple pathologies, including fatty liver disease, cirrhosis, hepatitis, infections and cancer (*10, 11*). To test if somatic mutations occur with a high frequency and accumulate rapidly with age we performed WGS on single primary hepatocytes from human donors varying in age between 5 months and 77 years of age. These cells were isolated from healthy human individuals shortly after death through perfusion of whole donor livers (Lonza Walkersville Inc.). Cell viability was higher than 80% and, after staining with Hoechst, hepatocytes were isolated via FACS into individual PCR tubes as diploid single cells (Fig. S1). In total we sequenced 4 single hepatocytes and bulk genomic liver DNA for each of eight human donors (Table S1). Each cell was subjected to our recently developed procedure for whole genome amplification (WGA) and sequencing (*5, 6*). Somatic single nucleotide variants (SNVs) in single cells were identified relative to bulk genomic DNAs at a depth of ≥ 20X using a combination of VarScan2, MuTect2 and Haplotypecaller software with certain modifications (Methods, Table S2). Overlapping mutations, revealed by all three analytical tools, were exclusively considered for further analysis.

After adjusting for genomic coverage, the number of SNVs per cell were found to vary between 357 and 5206 with one extreme outlier of 35640 excluded from the statistical model (Fig. 1A, Table S3). The number of mutations per cell was found to increase significantly with the age of the donor (*P*=3.18 x 10^−4^), with median values of 1139±776 SNVs per cell in the young group (≤30 years, n=15 cells), and 4533±898 SNVs per cell in the old group (≥51 years, n=16 cells) (Fig. 1A). The median number of mutations per cell in hepatocytes from the youngest donor were in the same range as what we recently reported for primary human fibroblasts from young donors, i.e., 1027 SNVs (5-month-old donor) and 926 SNVs (6-year-old donor) per cell, respectively (*5, 6*).

**Fig. 1.**
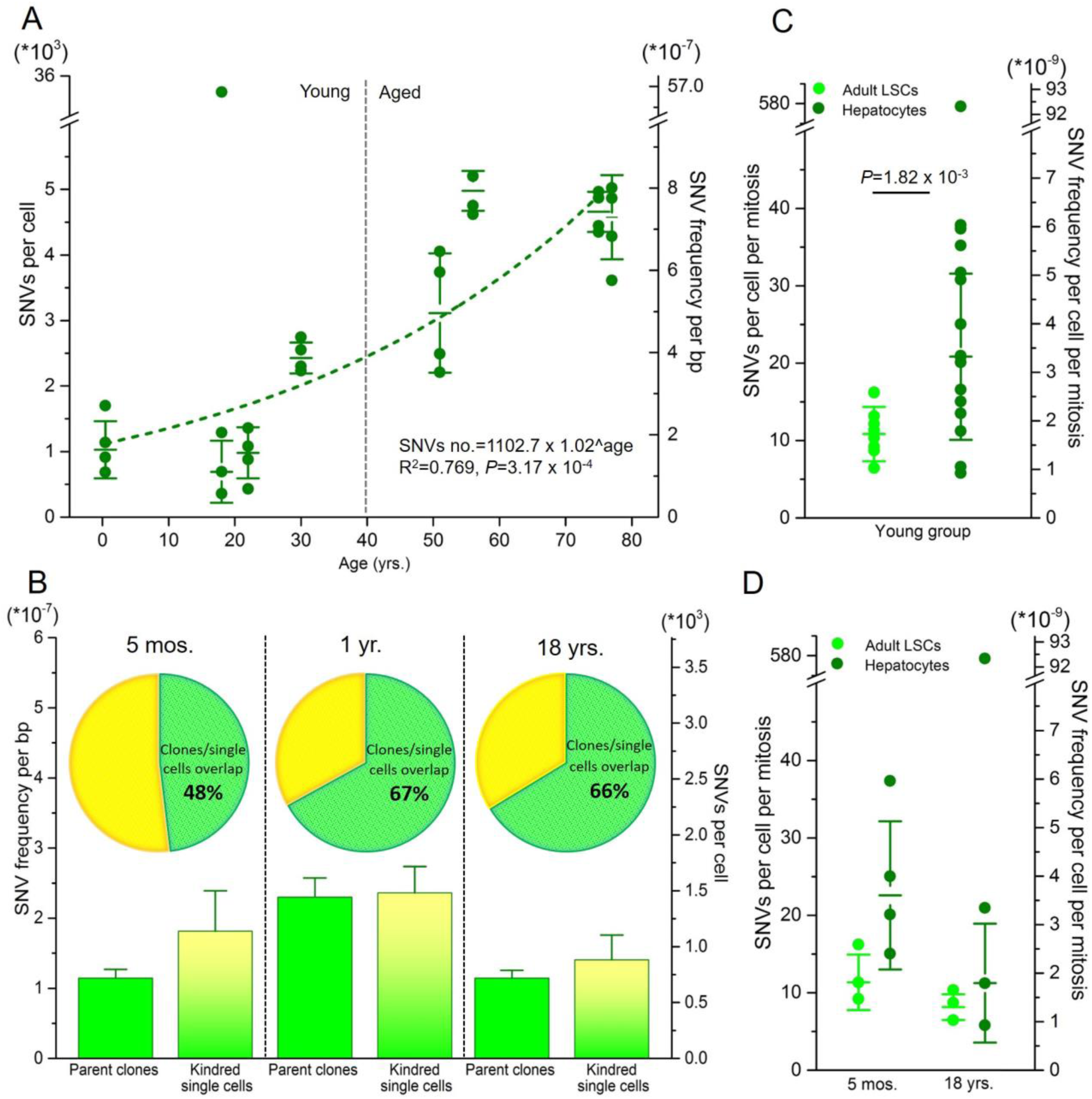
Somatic mutational levels in normal human liver cells. (A) SNV number per cell increases as a function of age in differentiated human hepatocytes. Dark green dots indicate individual cells, altogether 4 hepatocytes per each of 8 donors of different age were analyzed. The regression was performed on the median numbers of mutations of the 4 cells from each donor, using an exponential model excluding one outlier cell from the 18-year-old subject from the statistical model. (B) Mutation frequency in parent clones (light green) and their kindred cells, derived from adult liver stem cells (LSCs; gradient light green/yellow) of three young donors. The fraction circle indicates highly overlapping somatic SNVs, with shared SNVs identified in both parent clone and its kindred single cell indicated in green and SNVs found only in the kindred cell but not in the parent clone indicated in yellow. (C) Increased mutation frequencies in differentiated hepatocytes compared to adult LSCs within the young group of donors ≤30 yrs. Light green dots indicate LSCs single cells (n=10 from 3 donors); dark green dots indicate single hepatocytes (n=15 from 4 donors). Mutation frequencies were corrected for the estimated number of cell divisions, which is higher in the LSCs. (D) SNV frequency in LSCs and differentiated hepatocytes from the same subjects, corrected for estimated number of cell divisions.

Importantly, our results on mutation frequency in human hepatocytes from older donors seem in conflict with results obtained with stem cell-derived liver organoids as surrogates for single cells (*4*). In this study the number of mutations per total diploid genome, i.e., the genome of the stem cell that gave rise to the organoid, varied from 1109 SNVs in individuals of 30-year-old to 2302 SNVs in individuals of 55-year-old. This is about 2 times lower than the numbers we here report for individuals of the same age, i.e. from 2426 SNVs in 30-year-old donors to 4978 SNVs in 56-year-old donors (*P*=4.43 x 10^−4^, Fig. S2, Table S3) (*4*). It occurred to us that our data in cells from older subjects could be overestimates due to amplification artifacts, for example, due to excessive age-related DNA damage accumulation in liver. For our aforementioned previous studies on human primary fibroblasts we validated single-cell data by also analyzing unamplified DNA from clones derived from cells in the same population (*5*), similar to the organoids in the study described above (*4*). Here we generated liver-specific clones by plating the hepatocyte cell suspension in selective medium for stem cell expansion (Methods). Under these conditions the differentiated hepatocytes died within 5-7 days while the liver stem cells (LSCs) present in the population attached and were propagated without differentiating, which was confirmed using biomarker analysis (Methods; Fig. S3) (*12, 13*). In this way we managed to grow clones from liver cell populations from 3 young individuals: a 5-month old, a 18-year old and a 22-year old. In addition, we obtained from a commercial source one sample of human postnatal LSCs, which were expanded and also grown into clones in the same way.

Stem cell clones were analyzed from the 1-year old, the 5-month old and the 18-year old and subjected to WGS. From each clone we also obtained kindred single cells, which were processed for sequencing as described above. We then tested if all mutations called in the single cells were also present in their parent clone and most of them were (Fig. 1B). This is very similar to what we previously reported for human single fibroblasts and clones derived from the same population of cells (*5*), which underscores the validity of our single-cell mutation detection method, also in liver cells. The mutations found in the kindred single cells, but not in the clone are not likely to be artifacts, but instead either mutations missed during variant calling in the clone (not unlikely, due to our stringent filtering) or *de novo* mutations arising in the individual cells during clone culture and expansion. Of note, amplification artifacts due to DNA damage are unlikely to be called by our pipeline, which excludes most SNVs present in less than 25% of the reads, i.e., corresponding to one damaged strand out of four DNA strands (*6*).

We then tested directly if mutation frequency in the single cells derived from the clones, defined as LSCs, is lower than in differentiated hepatocytes. After correcting for the difference in the estimated number of cell divisions (Methods), the results show that somatic mutation frequencies were indeed significantly lower in the LSCs than in the differentiated hepatocytes (2-fold) from young donors, i.e., 11 SNVs vs. 21 SNVs per cell per mitosis, respectively (*P*=1.82 x 10^−3^, two-tailed student’s t-test) (Fig. 2C, Fig. S4A, Table S3). A reduced mutation rate in LSCs could explain the fairly modest age-related increase reported previously for stem cell-derived organoids (Fig. S4A) (*4*) and the much more robust increase we observe in single differentiated hepatocytes (Fig. 1A). The tendency of differentiated hepatocytes to accumulate mutations to a much higher level than stem cells is further confirmed by the significantly higher cell-to-cell variation among the former (Fig. 1C, D, *P*=2.86 x 10^−3^, Levene’s test). These observations are in keeping with the idea that stem cells preserve their genome integrity by remaining quiescent (minimizing replication-based errors) or through an enhanced capability to prevent or repair DNA damage (*14, 15*).

**Fig. 2.**
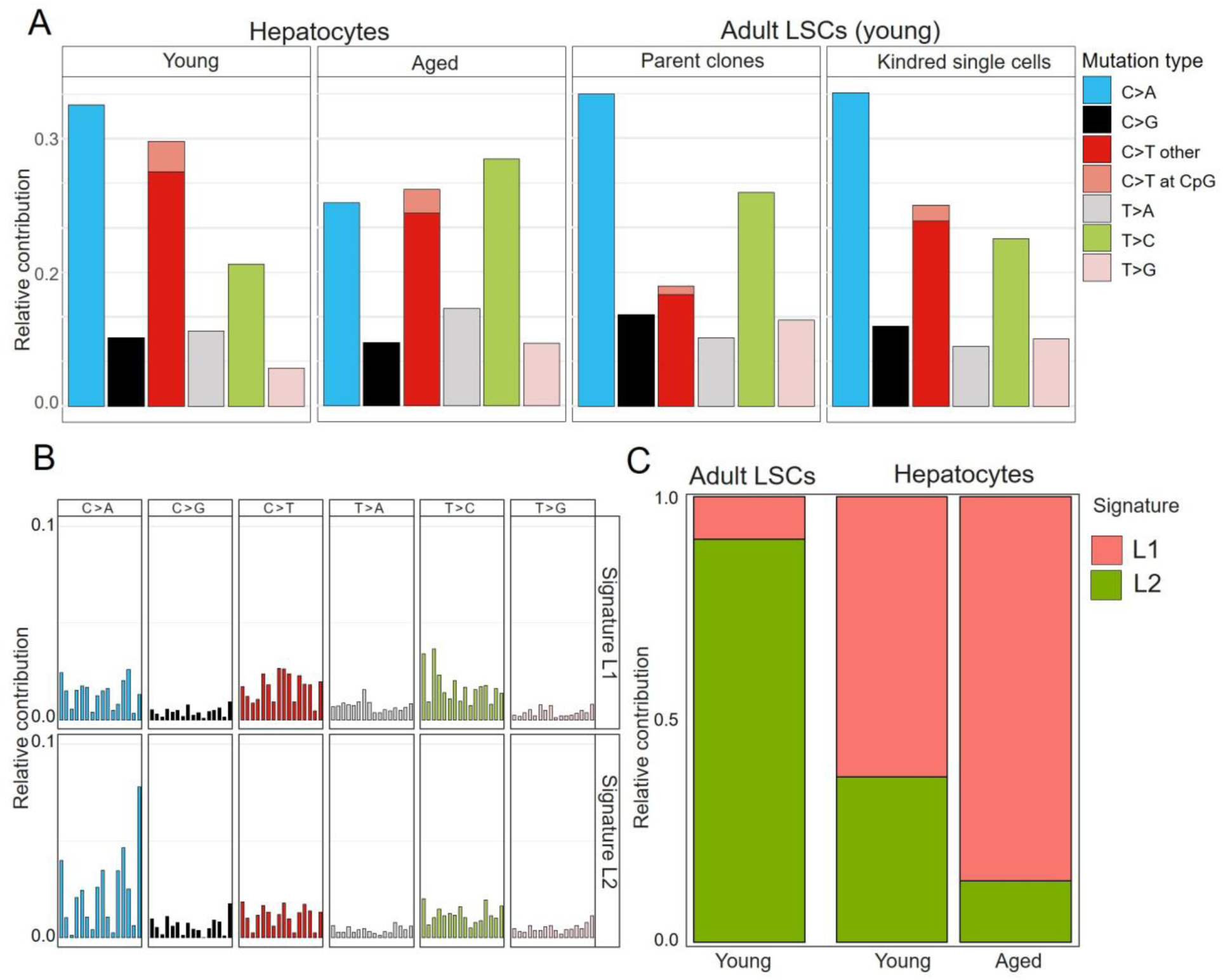
Mutational spectra in normal human liver cells. (A) Relative contribution of the indicated mutation types to the point mutation spectrum for each liver sample group. Data are represented as the mean relative contribution of each mutation type in sample groups of young and aged differentiated hepatocytes (15 cells from 4 donors ≤30 yrs and 16 cells from 4 donors ≥51 yrs) and adult LSC-derived parent clones and their kindred single cells. (B) Two mutational signatures (L1 and L2) identified by non-negative matrix factorization analysis of the somatic mutation collection observed in differentiated hepatocytes and LSCs. (C) Contributions of signatures L1 and L2 to all SNVs in young and aged hepatocytes and young LSCs.

Next, we analyzed the mutational spectra in LSCs and differentiated hepatocytes (Fig.

2A, Table S4). In differentiated hepatocytes the most common mutations were GC to AT transitions and GC to TA transversions (Fig. 2A; Fig. S5). These mutations are known to be induced by oxidative damage (*16*), which itself has often been considered as a main driver of aging and age-related diseases (*17*).

The most rapidly increased mutation type with age was the AT to GC transition (*P*=1.83 x 10^−6^). This mutation can be caused by mispairing of hydroxymethyluracil (5-hmU), another common oxidative DNA lesion. Alternatively, T>C lesions are commonly induced by mutagenic alkyl-DNA adducts formed as a result of thymine residue alkylation (*18, 19*). Notably, certain minor alkyl-pyrimidine derivatives can escape repair, accumulate during aging and lead to T>C mutations much later (*19, 20*). Mutation spectra of the LSCs and LSC clones revealed a lower fraction of GC to AT transitions than GC to TA transversions as compared to hepatocytes from the young group. It is possible that the quiescent state of LSCs and the absence of a functional role in liver metabolism results in a different ratio of these two main mutation types associated with reactive oxygen species. Interestingly, in the human LSCs derived from clones the relative frequency of the GC to AT transition mutations is again slightly, albeit significantly, increased as compared to the clones themselves (*P*=7.43 x 10^−4^; Fig. 2A). Kindred single LSCs, derived from parent LSC-clones, representing the original LSCs, have undergone multiple rounds of cell division with ample opportunity for replication errors, for example, as a consequence of ambient oxygen to which these cells have been inevitably exposed during subculture.

In order to analyze mutation spectra more precisely, we performed non-negative matrix factorization (Methods) and specified two different mutational signatures (signatures L1 and L2) within the 3 groups of human liver cells (Fig. 2B). In differentiated hepatocytes from the group of young subjects signature L1 is prevalent, increasing further in old subjects (Fig. 2C). Indeed, this signature highly correlates with the liver-specific and age-associated signature A dominating in human liver organoids in the aforementioned organoid study of mutation frequencies and spectra (Cosine similarity=0.92; Table S5) (*4*), as well as with the liver-specific cancer signature (Cosine similarity=0.882; Table S5) (*21, 22*).

Signature L2, with its increased level of oxidative GC>TA transversions, dominates the mutation spectrum of human LSCs (Fig. 2C). This signature L2 highly correlates with the so-called culture-associated proliferation signature C found generally in all *in vitro* propagated organoid types in the aforementioned organoid study (Cosine similarity=0.859; Table S5) (*4*). This signature possibly reflects the stem/progenitor-like origin of hepatocytes and is still present in differentiated hepatocytes of the young individuals (Fig. 2C).

Next, we analyzed the distribution of the somatic mutations in human liver cells across the various sequence features of the genome. After pooling all mutations of the 16 differentiated cells from aged and the 15 cells from young individuals the large majority of mutations distributed randomly across the genome in both groups (Fig. S6A, Table S6). Interestingly, the number of mutations in non-intergenic regions of the genome was found to be significantly lower in the LSC clones and single cells than in the differentiated hepatocytes from old or young subjects. (Fig. S6B, Table S6). This observation suggests that unlike mature, low-proliferating hepatocytes quiescent LSCs are able to prevent mutations from functionally relevant sequences.

To specifically test the possible functional impact of somatic mutations in human liver we compared mutation rates in the functional genome with that in the genome overall (Table S7, Fig. S6B). Mutations in the functional genome were defined as those occurring in the liver transcribed exome or its regulatory regions. To identify the transcribed liver exome we used available data on gene expression levels in 175 previously described liver samples (Genotype-Tissue Expression (GTEx) Consortium) (*23*). Regulatory regions were identified as promoters of active genes or open chromatin regions, e.g., transcription factor (TF) binding regions, identified by ATAC-seq (ENCODE) (*24*).

The normalized ratio of total to functional SNVs was found to be 1.6 in pooled adult LSCs as compared to a ratio of 1 in differentiated hepatocytes (*P*=5.34 x 10^−4^, Wilcoxon signed-rank test, two-tailed). This also is in keeping with increased protection against deleterious mutations in progenitor/stem cells. This ratio of 1 between total and functional SNVs remained stable in differentiated hepatocytes with age (*P*=0.779, Wilcoxon signed-rank test, two-tailed) (Fig. 3A, Table S8). This indicates that there is essentially no selection against deleterious somatic mutations in low-proliferating hepatocyte populations during aging.

**Fig. 3.**
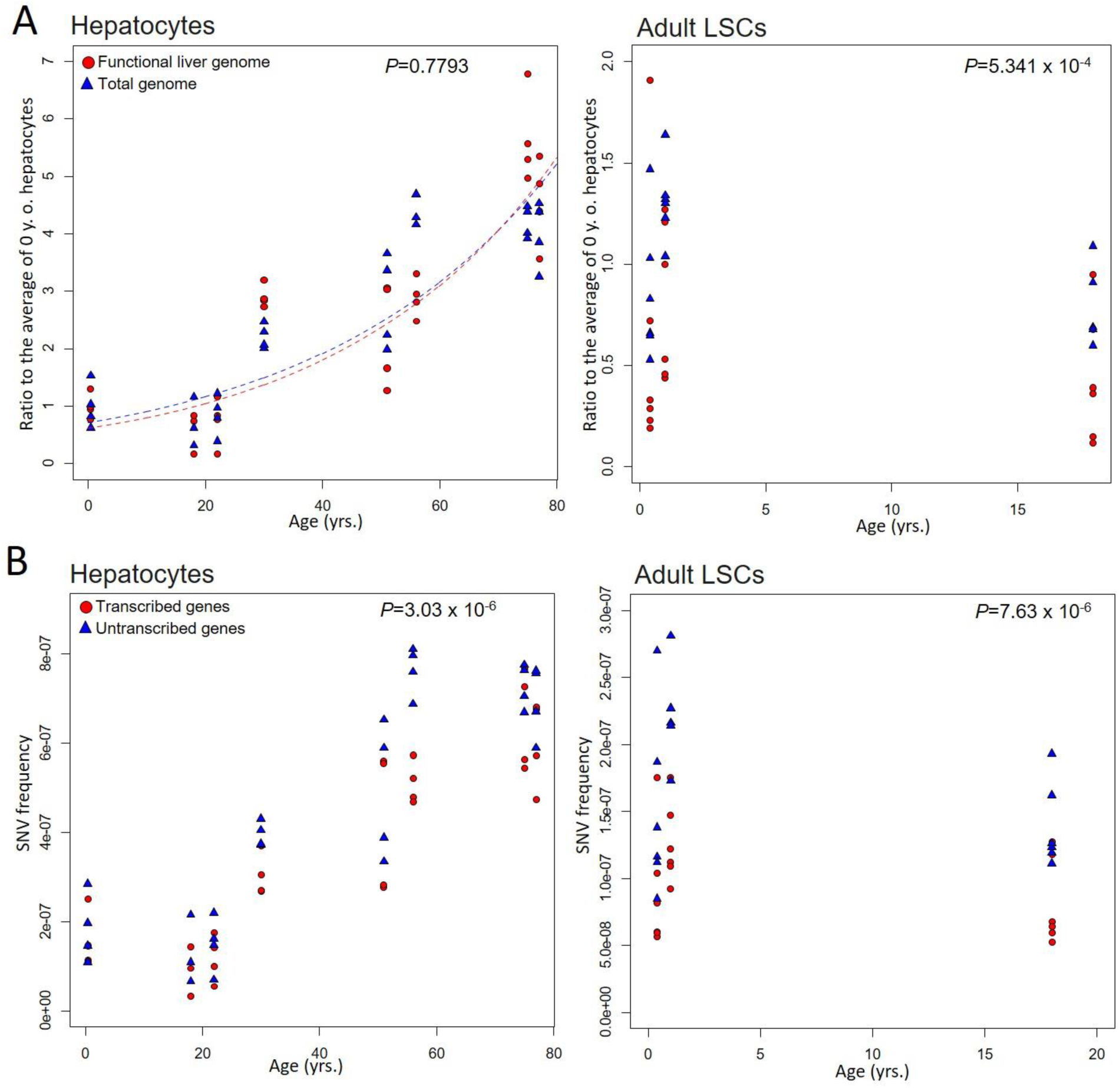
SNV levels in the functional genome and genome overall in normal human liver cells. (A) Each data point represents the number of mutations per cell. Mutations in the functional genome are shown in red and those in the genome overall in blue. Ratio of total vs. functional SNVs accumulated with age in differentiated hepatocytes remained stable indicating minimal selection pressure with aging. This ratio in LSCs is 1.6 times as high as in differentiated hepatocytes suggesting increased protection against deleterious mutations in progenitor/stem cells compared to functional terminally differentiated cells. (B) Number of SNVs affecting the transcribed part of the liver genome (red) is significantly lower than that in the non-transcribed part (blue) in both differentiated hepatocytes across all donor ages and LSCs.

Furthermore, we compared mutation rates in the transcribed and non-transcribed liver cell genome. Transcribed liver genome was defined as sequences with expression values ≥ 1 transcripts per kilobase million (TPM), while non-transcribed genome included all sequences with expression values < 1 TPM (GTEx) in liver tissue (*23*). The results indicated a significantly lower number of SNVs affecting the transcribed part of the liver genome than the non-transcribed part. This was found in differentiated hepatocytes across all donor ages (*P*=3.03 x 10^−6^, Wilcoxon signed-rank test, two-tailed) as well as in the liver stem cells/clones (*P*=7.63 x 10^−6^) (Fig. 3B, Table S8), suggesting active transcription-coupled repair in normal human liver (*25*).

In summary, using our advanced single-cell sequencing method we found that somatic SNV level in normal liver significantly increases with age, reaching as much as 4 times more mutations per cell in aged humans than in young individuals. Of note, the numbers of mutations in aged liver are much higher than what has previously been reported for aged human liver organoids (Fig. S2) (*4*) and also higher than recent results reported for aged human neurons (Fig. S7). Since we essentially ruled out that many of these mutations are artifacts of the amplification system the most likely cause of this high mutagenic activity in human liver is the high metabolic and detoxification activity in this organ. Importantly, somatic mutation levels in differentiated, functional human hepatocytes were found to be much higher than in LSCs. This means that clonal surrogates for cells in vivo do not always accurately represent the mutation loads of differentiated cells, which makes predictions of a functional impact of somatic mutations from such clonal data very difficult. While we do not know the mechanism(s) of reduced spontaneous mutation loads in stem as compared to differentiated cells, similar findings have been reported by others (*26, 27*) and it is possible that stem cells protect their genome integrity by remaining quiescent, thereby avoiding replication errors, and/or being equipped with superior DNA repair mechanisms as compared to their differentiated counterparts.

Finally, an important question is the possible functional impact of random somatic mutagenesis on the aging phenotype. While from our current data we cannot conclude direct cause and effect relationships, our observation that the functional part of the genome accumulated numerous mutations suggests that aging-related cellular degeneration and death could at least in part be due to somatic mutations.

## Acknowledgments

We thank the Flow Cytometry Core at the Albert Einstein College of Medicine for assistance in single cell sorting and collection.

## Author contributions

J.V. A.Y.M. and K.B. conceived this study and designed the experiments. O.M., M.K. and A.W.V. provided field-specific study expertise and logistics. K.B. performed the experiments. S.S. and X.D. analyzed the data. K.B., S.S. and J.V. wrote the manuscript.

Materials and Methods

Figures S1-S7

Tables S1-S8

References (*28–45*)

