## Supplementary text and figures for "Single-cell analysis reveals different age-related somatic mutation profiles between stem and differentiated cells in human liver"

**This PDF file includes:**

Materials and Methods
Figures S1 to S7
Captions for Tables S1 to S8

**Other Supplementary Materials for this manuscript include the following:**

Tables S1 to S8

### Materials and Methods

Human specimens

Frozen human hepatocyte samples were purchased from Lonza Walkersville Inc. All eight selected hepatocyte donors were healthy subjects of various age, gender and ethnicity (Table S1) without any liver cancer or other liver pathology history. These cells had been isolated using a gold-standard, two-step liver/liver lobe perfusion procedure. Cells were suspended in 2-5 ml of media, cells, counted with Trypan blue to estimate viability (higher than 80%) and frozen in DMSO/liquid nitrogen (https://www.lonza.com). One specimen of frozen human neonatal liver stem cells (LSC) from a one-year old donor was purchased from Kerafast, Inc. (https://www.kerafast.com). These cells had been derived by Sherley laboratory and characterized to confirm their stem cell identity (*28-30*).

Single hepatocyte collection

After thawing, hepatocyte suspensions were used to collect single hepatocytes into individual 0.2 ml PCR-tubes with 2.5 µl of PBS by means of fluorescent activated cell sorting (FACS; FACSAria, Becton Dickenson). Selection of the target hepatocyte population was based on the large cell size of hepatocytes (FSC/SSC parameters) along with the additional fluorescent staining for DNA content and cell viability. Briefly, bulk hepatocytes suspension samples were prior stained according to the manufacturer's protocol with the viable DNA-binding dye Hoechst 33342 (Life Technologies) to discriminate cells with a standard diploid chromosome set and LIVE/DEAD Cell Vitality Assay Kit $C_{12}$ Resazurin/SYTOX™ Green (Thermo Fisher Scientific) to select viable healthy cells. Typical FACS sorting lay-out is shown in Fig S1. Upon sorting, tubes with single cells were frozen on dry ice and kept at -80˚C until use.

Adult LSC polarization and culture

Neonatal LSCs from the one-year old donor (Kerafast, Inc.) were cultured in polarization media (DMEM, 10% dialyzed FBS (Invitrogen), 1.5mM Xanthosine (Sigma), 1x Penicillin/Streptomycin, 20ng/ml EGF human (Invitrogen), 0.5ng/ml TGF beta human recombinant (Sigma)) according to the manufacturer's protocol (Kerafast, Inc.) (*28-30*). These cells served as a control to characterize *de novo* isolated and polarized LSCs.

Additional LSC cultures were isolated and polarized and characterized from the bulk commercial hepatocyte suspensions (Lonza Walkersville Inc.) using previously described protocols with specific modifications (*12, 13*) combined with the aforementioned Kerafast protocol for neonatal LSCs. Briefly, bulk suspension hepatocytes (0.5-1 x $10^6$ of cells) were transferred to polarization media as described for the neonatal LSCs and cultured on cell-adhesive 12 well-plates for 3-4 days. After that all suspension non-attached hepatocytes were removed and fresh media was added to the remaining rare population of attached progenitor cells. After 1-2 weeks of culture and media changes attached cells symmetrically divided growing to mixed clonal populations of polarized adult LSCs. These cultures were frozen at early passage until further use. Phenotype of the polarized cells was analyzed and characterized for the presence of several specific surface stem cell and epithelial progenitor cell markers, e.g. EpCAM, Lgr5, CD90, CD29, CD105, CD73, upon staining with antibodies by means of multicolor flow cytometry analysis (LSRII, Becton Dickinson) as recommended previously (*12, 13, 31, 32*). Characteristic FACS profiles and specific phenotypes for commercial LSCs (control) and two manually isolated and polarized LSC lineages are shown in Fig. S3.

Adult LSC clones and single cell establishment

Single-cell derived parent clones and their kindred single cells were prepared and collected using CellRaft arrays (Cell Microsystems) as described previously (*5*). Briefly, LSC suspension was plated on a CellRaft array consisting of 12,000 individual portable rafts for single cells at the required density of 5000 cells per array. After 4-8 hours individual LSCs were elongated and attached to the array surface locating on individual rafts. After attachment, the medium with floating cells was replaced and single cell positions were marked and tracked during the following 7-10 days to detect dividing cells and growing individual single cell-derived clones. Once the colony/clone reached confluence on the raft (8-10 cells per raft), it was dislocated from the array with a positioned automatic needle and transferred with a magnetic wand to a 96-well plate. Upon reaching confluence, single-cell derived clones were trypsinized and subsequently transferred to 24-well plates, then 12-well plates, 6-well plates and, finally, 10 cm plates to reach a total amount of 1.5-3 x $10^6$ cells per each parent clone. Altogether, the process of establishing a clone from a single cell took about 25-30 days.

Individual single cells from the parent clones were collected, also using CellRafts, and transferred to a 0.2 lµ PCR tube containing 2.5 l µPBS. Presence of a single raft was observed under a magnifying glass. Upon single-cell collection, tubes were fast frozen on dry ice and kept on -80˚C until further use.

Single cell whole genome amplification

Single hepatocytes from each individual subject were subjected to whole genome amplification (WGA) using our modified procedure of low temperature cell lysis and DNA denaturation followed by multiple displacement amplification (MDA) as described (*5*). As positive and negative controls for WGA we used 1 ng of human genomic DNA and DNA-free PBS solution, respectively. Resultant MDA products were purified using AMPureXP-beads (Beckman Coulter), amplified DNA concentration was measured with Qubit High Sensitivity dsDNA kit (Invitrogen Life Sciences). To verify sufficient and uniformly amplified single cell MDA products we performed the 8-target locus-dropout test as described previously (*5*). Selected confirmed samples (4 single cell MDA products per each individual subject) were further subjected to library preparation and whole-genome sequencing.

Genomic DNA and clone-derived DNA extraction

Human bulk genomic DNA was collected from total cell suspensions (for hepatocytes and respective polarized LSC) using DNeasy Blood & Tissue Kit (Qiagen) according to the manufacturer's protocol. LSC clone-derived DNA was extracted from clones of at least 1.5-2.5 x $10^6$ cells in a similar way. DNA concentration was quantified with Qubit High Sensitivity dsDNA kit (Invitrogen Life Sciences) and DNA quality was controlled by 1% agarose gel electrophoresis.

Library preparation and whole genome sequencing

The libraries for Illumina next-generation WGS were generated from 0.2-0.4 µg genomic DNA, clone-derived bulk DNA and single cell MDA DNA human samples using NEBNext Ultra II FS DNA Library Prep Kit for Illumina (New England Biolabs). The libraries were sequenced with 2 x 350 bp paired-end reads on an Illumina HiSeq X Ten sequencing platform by Novogene, Inc.

Next generation WGS at a minimal depth of 20 X base coverage was performed on four individual mature hepatocytes per each human subject (8 human subjects, 32 single cells in total) (Table S2). Bulk DNA from two or three LSC-derived clones and MDA products from 3-4 corresponding kindred single cells per donor (3 donors, 8 parent clones and 10 kindred single LSCs) were sequenced similarly.

Alignment for whole-genome sequencing

For all samples, adapter and low-quality reads were trimmed by Trim Galore (version 0.3.7). Quality checks were performed before and after read trimming by FastQC (version 0.11.4). The trimmed reads were aligned to the human reference genome (GRCh37 with decoy) by BWA mem (version 0.7.10) (*33*). Duplications were removed using samtools (version 0.1.19) (*34*). The known indels and SNPs were collected from the 1000 Genomes Project (phase I) and dbSNP (build 144). Then, the reads around known indels were local-realigned, and their base quality scores were recalibrated based on known indels and single nucleotide variants, both via Genome Analysis Toolkit (GATK, version 3.5.0) (*35*).

Calling somatic single-nucleotide variants

We identified somatic mutations between each single cell and the corresponding bulk, and between each clone and corresponding bulk using three different variant callers: Varscan2 (*36*), Mutect2 (*37*) and HaplotypeCaller (*35*). To obtain high-quality mutation calls and avoid high false-positive rates in individual callers, we applied a comprehensive procedure in filtering. First, we only considered mutations on autosomes. Then, we considered mutations with a GATK phred-scaled quality score of at least 30 and excluded mutations overlapping with known SNPs from dbSNP. Furthermore, we required a minimum base depth of 20X and filtered mutations with variant-supporting reads in bulk. Moreover, we excluded mutations present in at least two cells in each individual to further exclude potential germline mutations. The mutations present in all three variant callers were considered as high confidence and collected. Finally, considering that amplification errors and/or non-uniform coverage could induce false-positive mutations in no more than 1/8 of the reads, we employed a binomial distribution to filter these potential false-positive mutations, which excluded most mutations present in 25% of the reads or less. To further check the power of our pipeline in filtering amplification errors, we also called the somatic mutations using alternative pipeline LiRA (Linked Read Analysis) (*38*) and compared our results with respective published neuron data (*8*) in Fig. S7.

Estimating mutation frequencies

Mutation frequencies were estimated by dividing the number of identified mutations by two times surveyed genome area due to diploid genome sequencing. The surveyed area per single cell/clone was calculated as the number of nucleotides with read mapping quality $\geq 20$ and position coverage $\geq 20$X. The absolute *de novo* mutation frequencies were also corrected for the number of cell divisions undergone by each cell with respect to liver cells turnover rate, donor age and cell culture period (Table S3), in order to eliminate SNVs accumulated as a consequence of replication errors. As the number of developmental mitoses, we used 45.1, as estimated previously (*7*). The turnover rate of human liver cells was reported to be approximately once per year (*39, 40*), which was used to calculate the number of divisions after birth depending on donor age. Finally, we added the additional number of cell divisions in LSCs during clone culture and expansion.

Detecting overlap between clone and kindred cell

For each mutation called in a kindred single cell, we examined variant-supporting reads in the corresponding clone (minimum base depth of 20X and mapping quality of 20) as described previously (*5*). The mutations found in the kindred single cells, but not in the clone are not likely to be artifacts, but instead either mutations missed during variant calling in the clone (not unlikely due to stringent filtering) or *de novo* mutations arising in the individual cells during clone culture and expansion. Mutations found in kindred single cell and appearing in at least 1 read in the parent clone were considered as the overlapping genotype. When there were no variant-supporting reads in the clone, the mutation was determined as kindred cell-specific. This assignment left some mutations with an unknown status more likely to be *de novo* mutations arising in the individual cells during clone culture and expansion. Amplification artifacts due to DNA damage are unlikely to be called by the used pipeline, filtering out all SNVs present in less than 25% of the reads (*6*).

Identifying mutation signatures

We pooled identified mutations in all individuals into three groups: LSC cells/clones, young liver, and aged liver. The integrated spectra of 6 mutation types in each group were plotted using the R package "MutationalPatterns" (*41*). Using NMF decomposition in the same package, we identified 2 signatures in normal human liver cells. To identify the potential origin of the mutational spectra, we compared our newly revealed signatures to published signatures associated with liver-specific organoids and various cancer tissues. Three tissue-specific organoid signatures were obtained from a recent study (*4*); thirty cancer mutation signatures were downloaded from the COSMIC database (http://cancer.sanger.ac.uk/cancergenome/assets/) (*22*) and original cancer signatures (*21*). The cosine similarity between newly identified and published signatures was calculated for comparisons (Table S5).

Annotation of functional genomes

All reported mutations were annotated based on the gene definitions of GRCh37.87. Mutations were further extracted from the functional genome, including transcribed genes, promoters and open chromatin regions. The nonsynonymous and synonymous mutations were identified by ANNOVA (*42*), while damaging and tolerated mutations were checked by SIFT (*43*) and PROVEAN (*44*). When damaging (SIFT) or Deleterious (PROVEAN), the mutation was marked as damaging, and when tolerated (SIFT) and Neutral (PROVEAN), a tolerated mutation.

The open chromatin regions were identified by ENCODE transcription factor binding regions in whole genome and ATAC sequencing data in the functional genome. Raw ATAC sequencing data were downloading from ENCODE (Experiment name: ENCSR373TDL) (*24*). The adapter and low quality ATAC sequencing reads were filtered using Trim Galore (version 0.3.7). Clean reads were aligned to the human reference genome (GRCh37) with Bowtie2 (version 2.2.3; option: -X 2000). Duplicated reads were removed with the Picard tool (version 1.119). Open chromatin regions were determined by MACS2 (version 2.1.1; option: callpeak -g hs --nomodel --shift -100 --extsize 200) (*45*).

Gene expression levels for liver tissue were obtained from GTEx (https://gtexportal.org/) (*23*). We defined the transcribed genes as those with expression level ≥ 1 TPM in all samples. Also, we separated the transcribed and non-transcribed genome by TPM ≥ 1 and <1 in all samples, respectively.

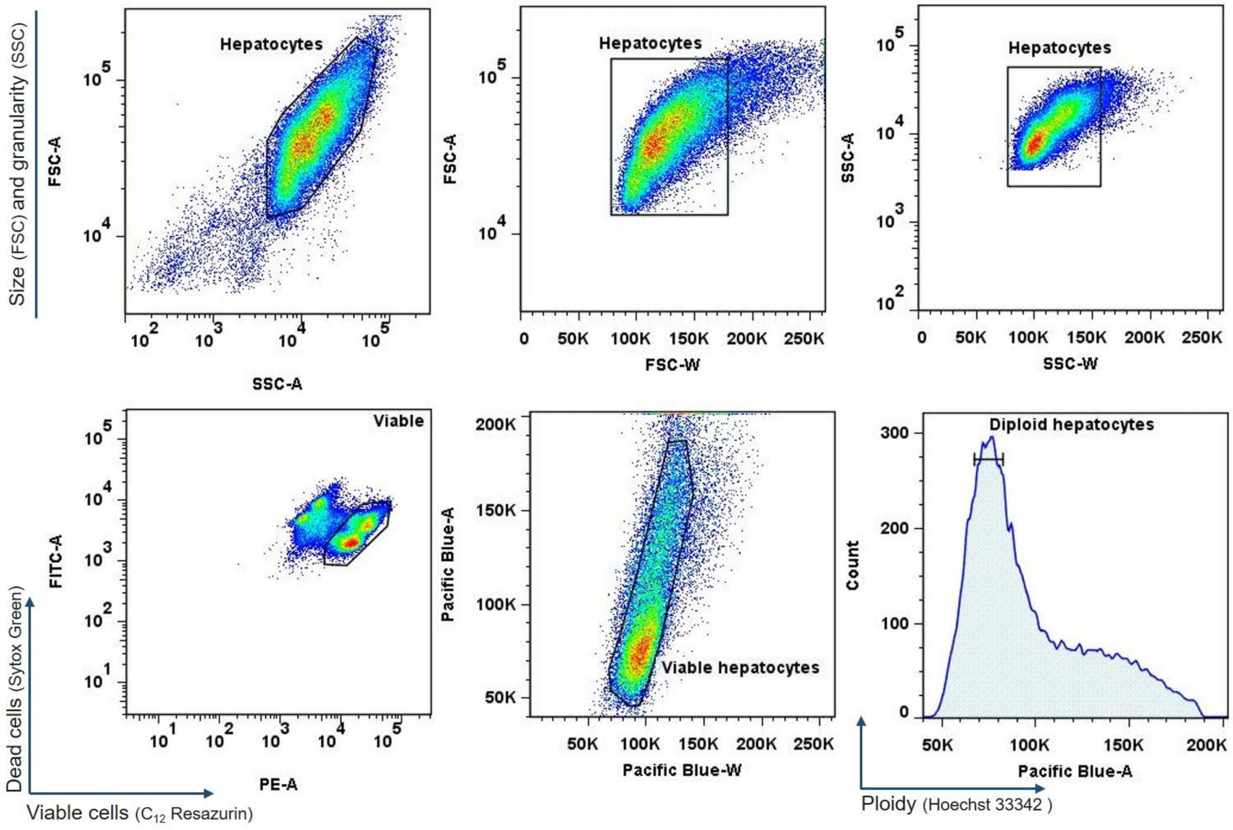


**Fig. S1.**

Typical FACS sorting layout to selectively discriminate and collect viable diploid human hepatocytes. The example shown is for normal hepatocytes from a 77 y. o. donor. Target population was selected based on large cell size (FSC/SSC parameters), DNA content (Hoechst 33342) and cell viability ($C_{12}$ Resazurin/Sytox Green).

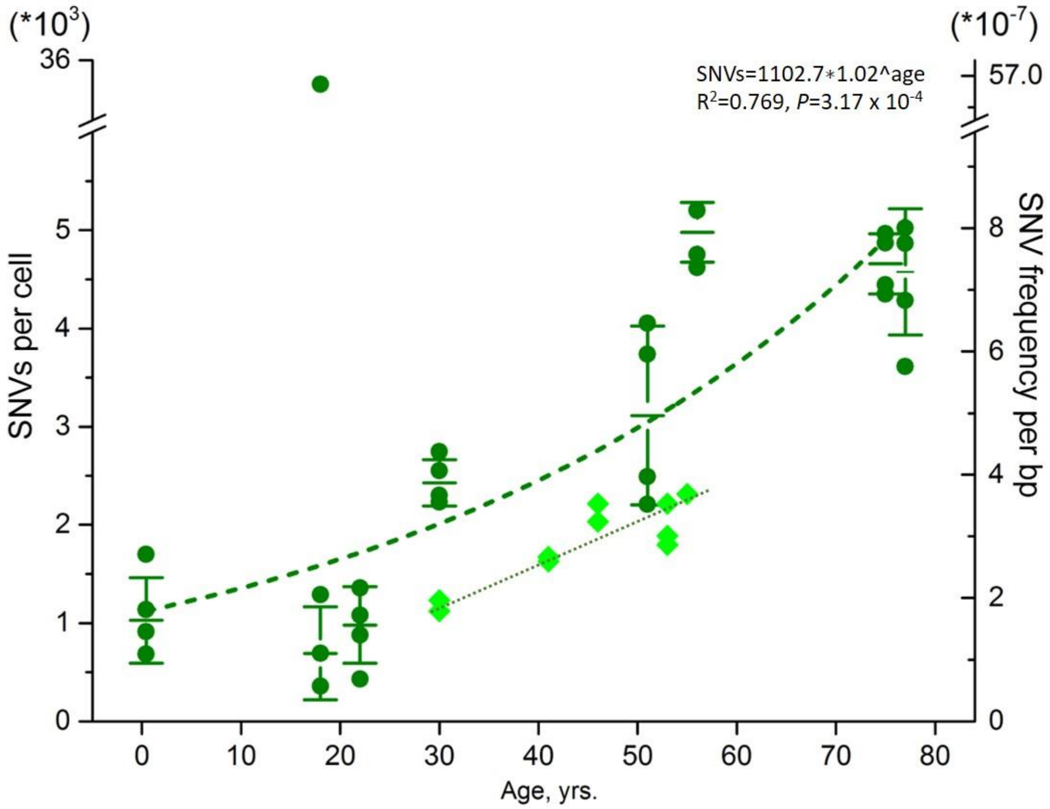


**Fig. S2.**

Comparison of SNV number and frequency in human differentiated hepatocytes (dark green circles) and adult liver stem cell-derived organoids (light green diamonds) from human donors of various age (*4*).

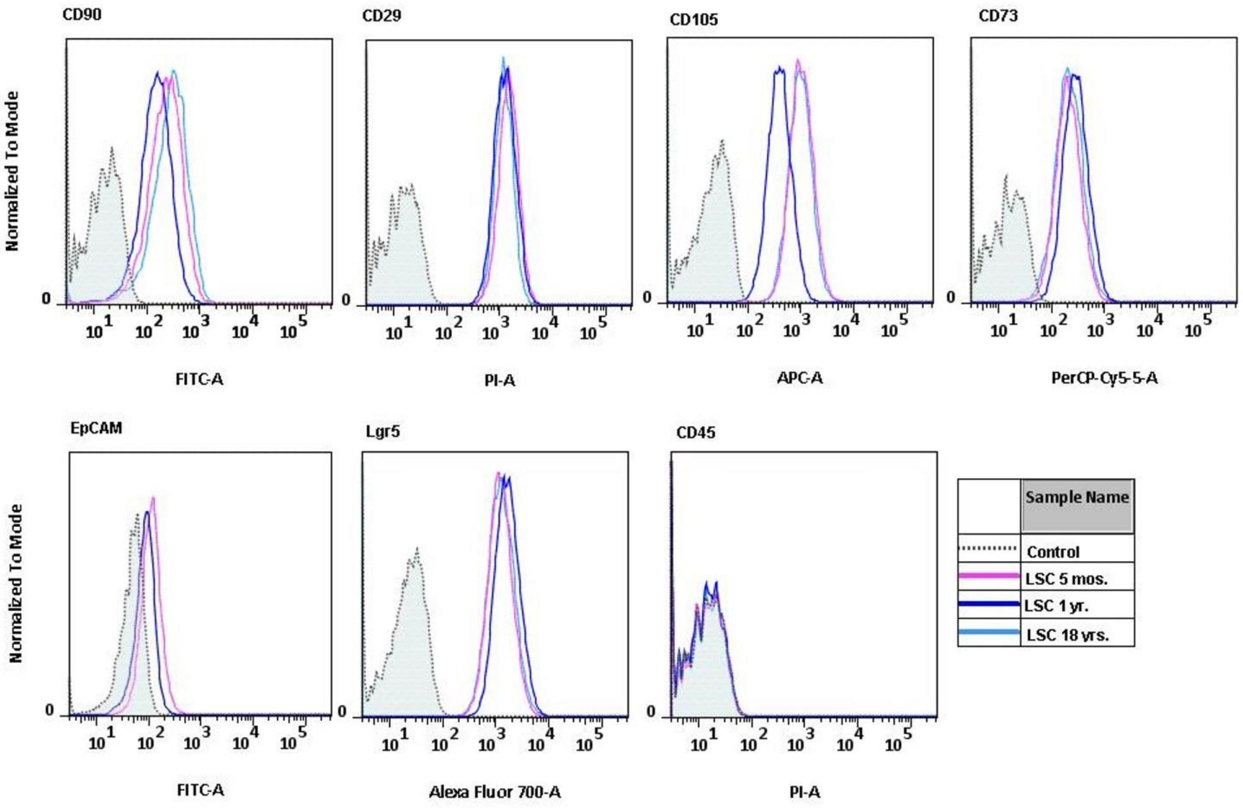


**Fig. S3.**

Phenotypic characterization of stem cell-specific markers (EpCAM, Lgr5, CD90, CD29, CD105, CD73) and negative markers (CD45) in control neonatal LSCs (LSC 1 yr., Kerafast, Inc.) and adult LSC populations (LSC 5 mos., LSC 18 yrs.) polarized and expanded from perfused hepatocyte cell fractions (Lonza Walkersville Inc.).

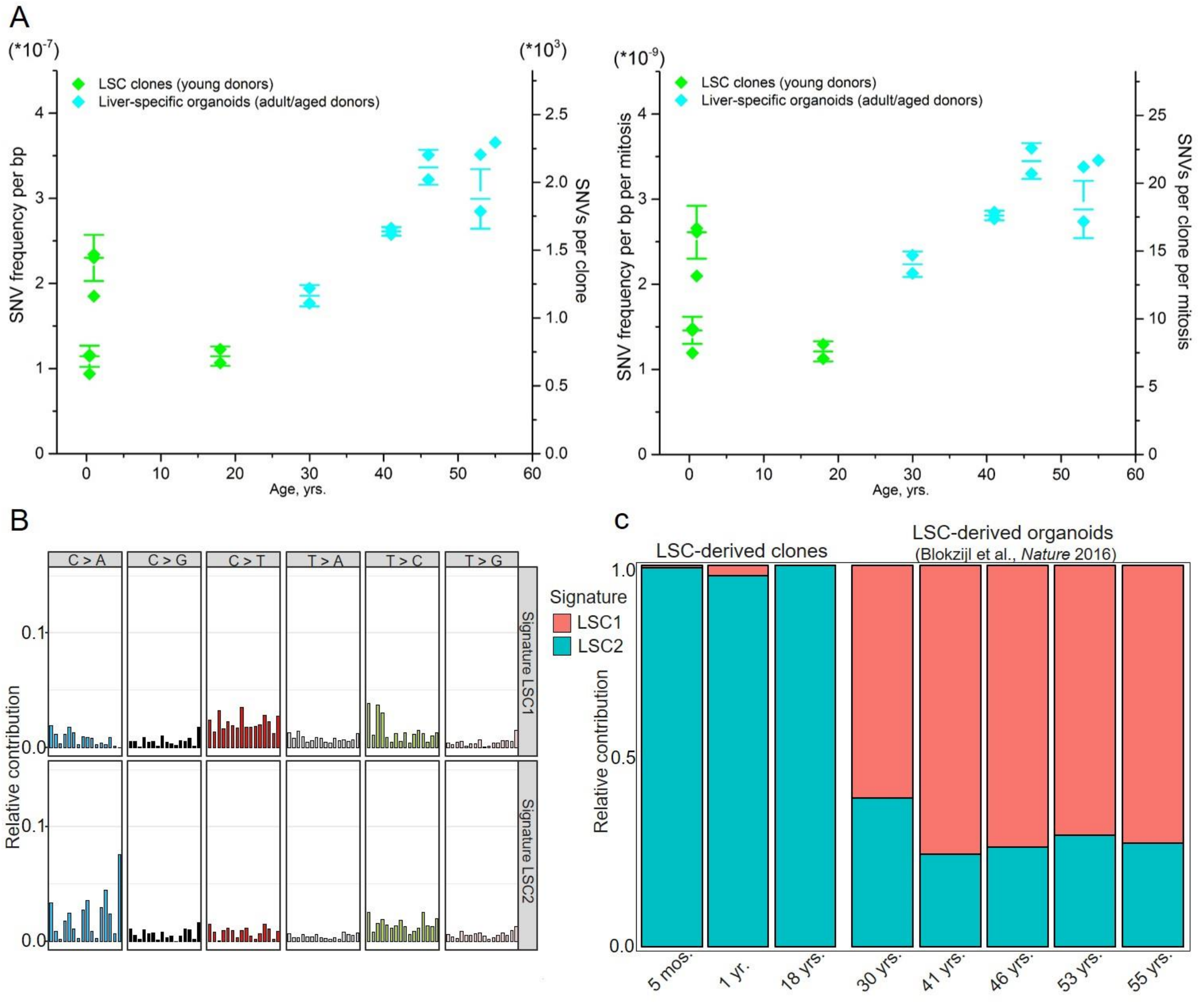


Fig. S4.
Comparison of adult LSC clones form three young donors (LSC 5 mos., LSC 1 yr., LSC 18 yrs.; light green diamonds) with liver-specific organoids from adult and aged donors (30-55 yrs.; light blue diamonds) (*4*). (A) SNV numbers and frequencies before and after correction for number of cell divisions with respect do donor age and clone/organoid expansion in culture. (B) Two mutational signatures (LSC1 and LSC2) identified by non-negative matrix factorization analysis of the somatic mutation collection observed in young adult LSC clones and adult/aged liver-specific organoids. (C) Contributions of liver-specific signature LSC1 and aging liver signature LSC2 to all SNVs in young adult LSC clones and adult/aged liver-specific organoids.

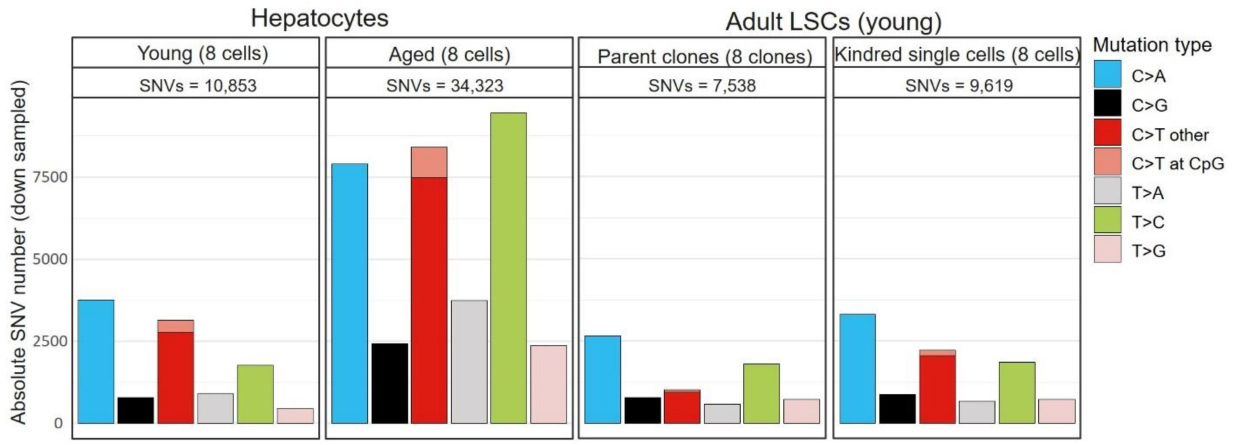


**Fig. S5.**

Absolute number of each of 6 indicated mutation types in each liver sample group. Raw total numbers of 6 mutation types within each group were down-sampled to the same number of samples (8 cells/clones per each group extracted randomly).

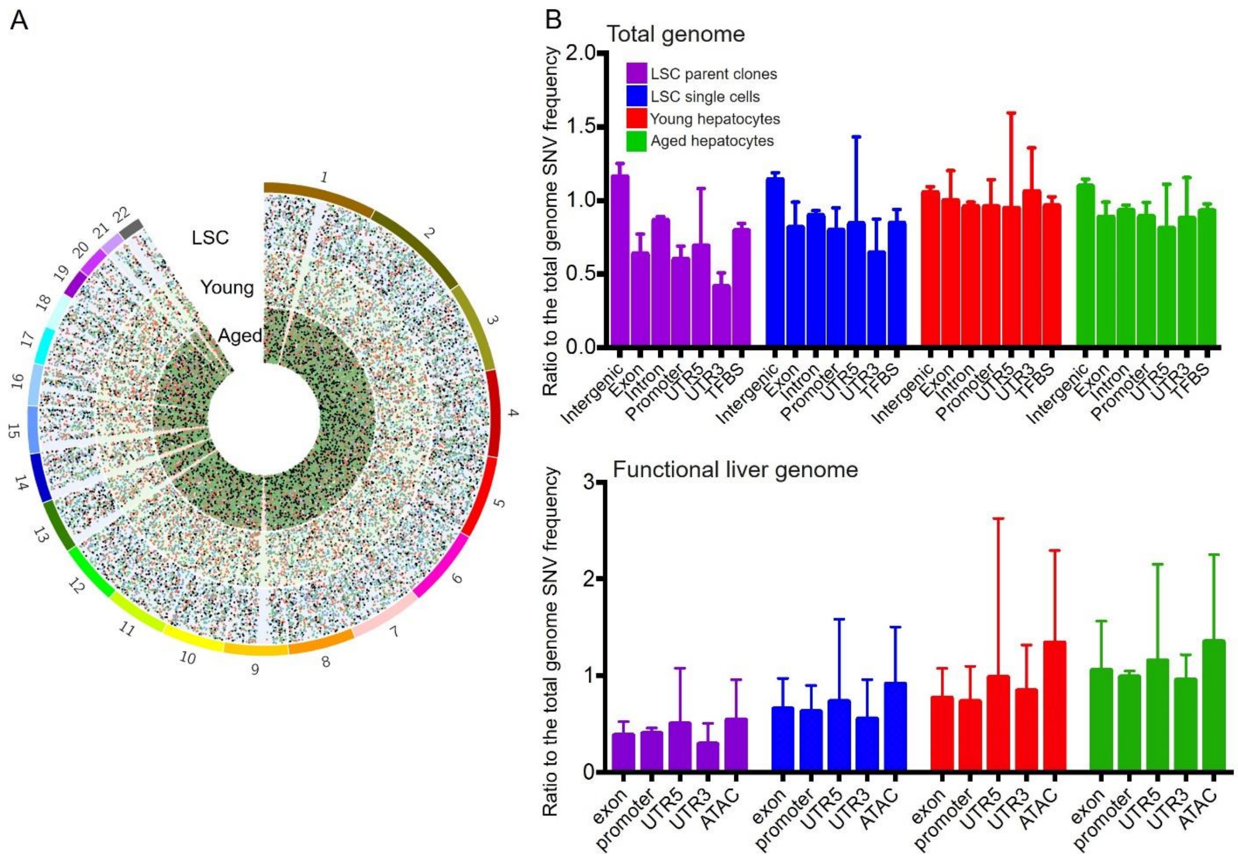


**Fig. S6.**

SNV distribution across total and functional liver genome within defined groups of pooled young and aged hepatocytes and young LSC parent clones and kindred single cells. (A) Circos diagram of random SNV distribution throughout the total genome in three groups of pooled LSCs, young and aged hepatocytes. (B) SNV distribution across specific genomic regions of total and functional genome in three groups of pooled LSCs, young and aged hepatocytes. Reduced SNV number in non-intergenic regions of the genome in the LSC clones compared to differentiated hepatocytes.

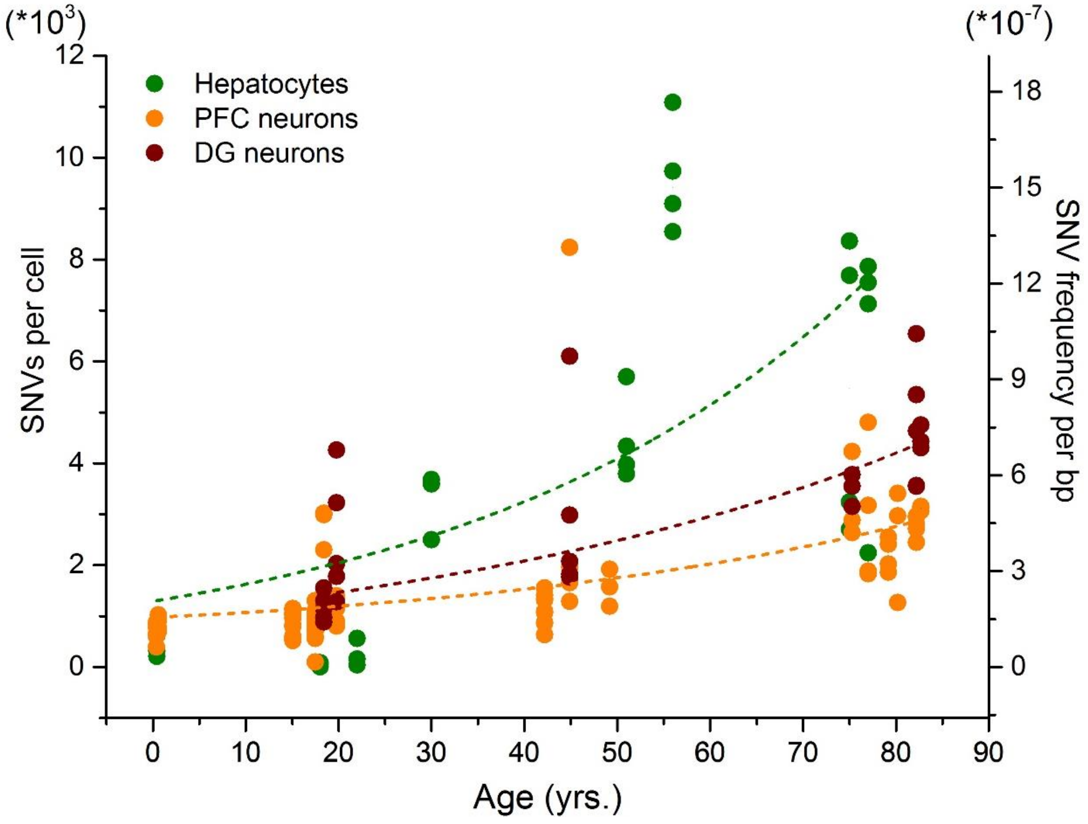


**Fig. S7.**

Comparison of SNV levels in differentiated hepatocytes and neurons (prefrontal cortex (PFC) and hippocampal dentate gyrus (DG)) identified using alternative pipeline LiRA (Linked Read Analysis) (*8*, *9*).

**Table S1. (separate file)**
Human liver donor information list.

**Table S2. (separate file)**
Whole-genome sequencing coverage and raw data on human liver cells.

**Table S3. (separate file)**
Final whole-genome sequencing data and mutation calling results on human liver cells.

**Table S4. (separate file)**
Distribution of 6 types of SNVs in total mutational spectra of human liver cells.

**Table S5. (separate file)**
Correlation of spectral signatures identified in human liver cells (L1 and L2 for hepatocytes vs. LSCs and LSC1 and LSC2 for LSC cells/clones vs. liver organoids) with cancer-related signatures and organoid-specific signatures.

**Table S6. (separate file)**
SNV frequencies and ratio calculated for specific genome sequences across total and functional liver genomes in human liver cells.

**Table S7. (separate file)**
Average number of SNVs per cell in indicated groups of pooled human liver cells distributed across total and functional liver genome within specific genome sequences.

**Table S8. (separate file)**
Summary of SNV frequencies in total, functional, transcribed and untranscribed genomes of human liver cells.
